# Single-cell multiome analysis supports α-to-β transdifferentiation in human pancreas

**DOI:** 10.1101/2025.02.14.638309

**Authors:** Michelle Y. Y. Lee, Omar Guessoum, Hilana El-Mekkoussi, Mitchell Conery, Elisabetta Manduchi, Jonathan Schug, Hélène Descamps, Deeksha Lahori, Tong Da, Chengyang Liu, Ali Naji, Benjamin F. Voight, Mingyao Li, Klaus H. Kaestner

## Abstract

Spontaneous transdifferentiation of pancreatic glucagon-producing alpha to insulin-secreting beta-cells has been observed in mouse but not in human islets^1^. Here, we analyzed the largest single-cell dataset of human islets to date, composed of 650,000 cells across 121 deceased organ donors, in search of transitional cell states. By integrating single-cell RNA-seq, single-nucleus ATAC-seq and single-nucleus multiome (joint RNA and ATAC profiling) datasets generated by the Human Pancreas Analysis Program (HPAP)^2,3^ we identified two previously undescribed cell populations (c11 and c13 cells), which together represent transitional states between alpha- and beta-cells. Some c11 cells are insulin-positive while others are glucagon positive, but none are double-positive. C11 cells repress alpha-cell identity genes and activate beta-cell specific genes. Moreover, the transcriptomic and epigenetic profiles of c11 and c13 cells indicate a transitioning phenotype driven by lineage-specific transcription factors. Genetic lineage tracing in primary human islet cells confirmed alpha-to-beta cell transdifferentiation. C11 and c13 cells exist in all islet samples regardless of disease statuses, with type 2 diabetic samples having significantly more transitioning cells than matched non-diabetic controls. The discovery of these transitional cell types suggests a possibility for future therapy – transdifferentiating alpha-cells to beta-cell through activation of the c11 gene program.

## Main

Pancreatic islet cell heterogeneity remains a topic of active debate, with numerous studies describing islet-cell dedifferentiation, transdifferentiation, or progenitor potential^4,5^. Interest in this field is driven by the limited understanding of the underlying causes of beta-cell loss and dysfunction in T2D, and the as yet unrealized potential to restore functional beta-cell mass as a treatment for the disease. Multiple studies in mice, where genetic lineage tracing and manipulation of key lineage-specific transcription factors is easily performed, have provided prior evidence for plasticity among mammalian pancreatic endocrine cells; for example, overexpression of the homeodomain transcription factor Pdx1 in alpha cells is sufficient to convert them to a beta-cell-like fate^6^, while ectopic activation of the alpha cell transcription factor Arx in beta cells causes activation of glucagon gene expression^7^. In addition, under conditions of extreme beta cell loss and fulminant diabetes, a small subset of alpha cells transition to an insulin-producing state^8^. Even under homeostatic conditions, there is a small but measurable conversion of alpha to beta cells in mice, as shown by careful genetic lineage tracing studies^1^. This degree of plasticity is plausible given that alpha and beta cells share a similar developmental lineage, diverging only at the final stage of differentiation, despite their opposing physiological roles. Interesting, human alpha cells have bivalent epigenetic marks at many beta-cell specific genes, suggesting they may be in a poised state for potential conversion into beta cells^9^. Nevertheless, spontaneous alpha-to-beta transdifferentiation has yet to be documented in human islets.

### Extensive single-cell data reveal two novel pancreatic endocrine cell populations in the human pancreas

Through efforts of the Human Pancreas Analysis Program (HPAP)^2,3,10,11^, we were able to analyze single-cell datasets from pancreatic islets consisting of 40 multiome, 85 scRNA-seq, and 62 snATAC-seq samples from four groups of deceased organ donors: nondiabetic (ND), autoantibody positive (AAB) but non-diabetic, type 1 diabetic (T1D), and type 2 diabetic (T2D).

Some of the samples were analyzed with multiple modalities; in total, there were 121 donors, with data from ∼650,000 cells (Fig.1a). We employed the multiome dataset as the ‘ground truth’ for cell type identification as it resolves cellular heterogeneity through both gene expression and chromatin accessibility profiles (Fig.1b), observing all sample groups across clusters (Fig. 1c). Following the recommended workflow from a recent benchmark study^12^, we mapped the single-modality datasets onto this multiome reference (Fig.1d-e). Across all three data types, we identified 11 canonical cell types showing expression of and chromatin accessibility at known marker genes (Supplementary Figs. 1 and 2), consistent with prior human islet single-cell studies^10,13^. However, two previously uncharacterized cell populations emerged, which we termed cluster 11 (c11) and cluster 13 (c13). In the multiome UMAP plot (Fig. 1b), cells belonging to these clusters were found in a continuum with alpha cells. When analyzing the RNA and ATAC modalities of the multiome data separately, c11 and c13 cells also separated out as individual clusters (Supplementary Fig. 1a-b). Similarly, when analyzing the multiome data from each donor individually (data not shown), c11 and c13 were consistently distinct from alpha cells. We observed c11 and c13 cells across all disease groups (Fig.1c, Supplementary Fig. 1c).

**Figure 1:**
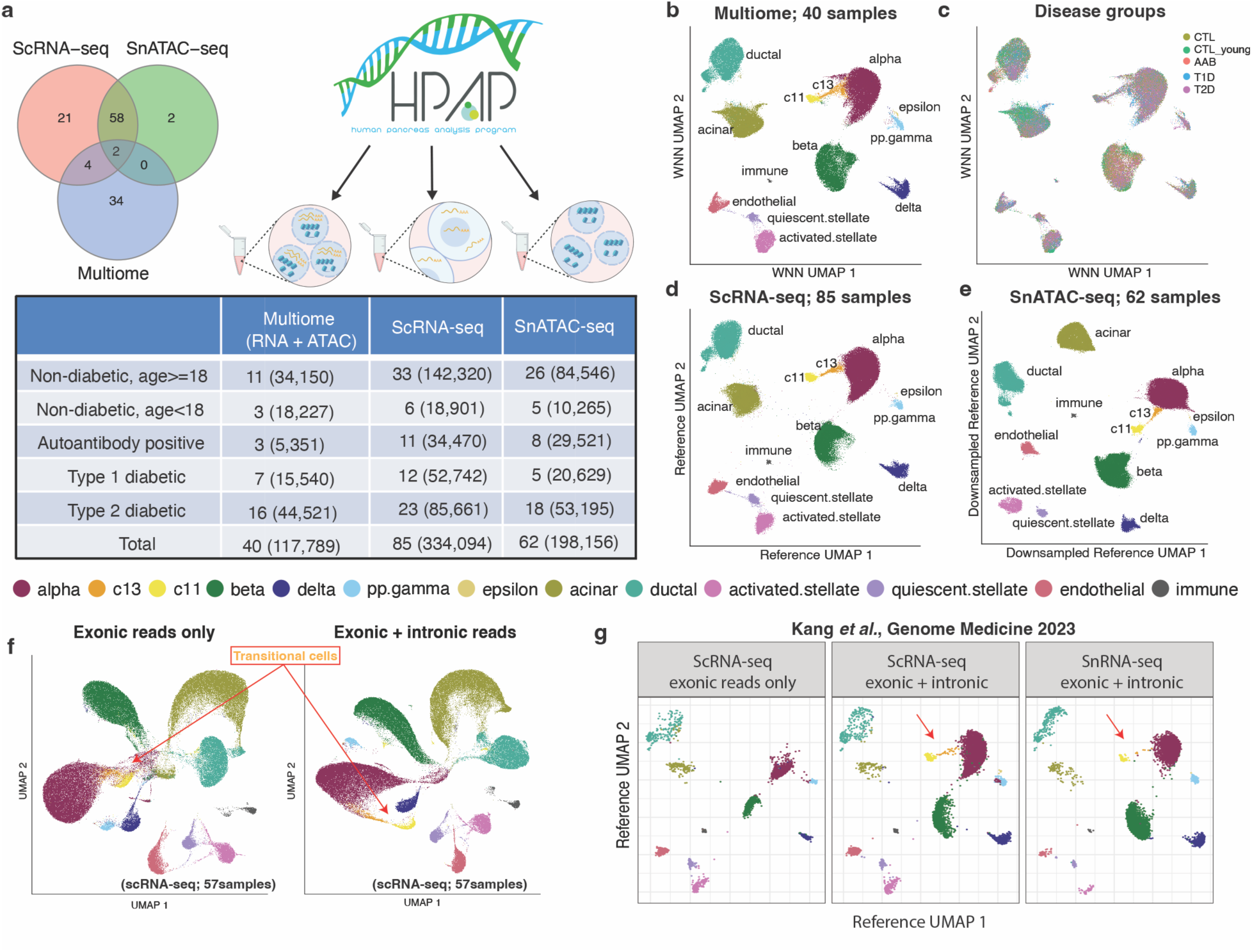
Transitional endocrine cell type identified in single-cell multiome datasets of the Human Pancreas Analysis Program (HPAP). (a) HPAP islet samples analyzed, showing the number of cells (in parentheses) by condition and methodology. The Venn diagram illustrates the overlap of donors across different data types. (b) UMAP visualization of 117,789 human pancreatic islet cells from the multiome dataset, generated using a weighted nearest neighbor (WNN) approach, which integrates batch-corrected RNA-seq and ATAC-seq data, processed via Seurat. (c) UMAP plot of the multiome dataset, colored by disease status. (d) UMAP plot of HPAP scRNA-seq cells projected onto the reference UMAP space of the multiome dataset. (e) UMAP plot of HPAP snATAC-seq cells, mapped onto a reference UMAP space created by down-sampling of the multiome dataset to 1,000 cells per donor. (f) UMAP plots generated from 57 scRNA-seq samples: (left) UMAP based on gene expression matrix considering only reads mapped to exonic regions; (right) UMAP using the same cells but including reads mapped to intronic regions. Transitional cells form a distinct cluster after incorporating intronic reads. (g) Re-analysis of the Kang et al. dataset^15^, with cells mapped to the HPAP multiome reference dataset. Transitional cells are identifiable in both snRNA-seq and scRNA-seq datasets only when intronic reads are included.

### Inclusion of intronic reads enables detection of novel transitional cell type

The question arose as to why the c11 and c13 clusters had not been discerned in prior scRNA-seq studies, including our own^10,14^. The answer emerged when we realized that most scRNA-seq workflows exclude reads mapped to the intronic regions of the genome; for instance, this is the default setting for Cell Ranger versions prior to 7.0.0. In contrast, the multiome gene expression data, which are derived from single-nucleus RNA-seq, include both intronic and exonic reads when analyzed by the 10x Genomics’ default analysis pipeline.

To determine if exclusion of intronic reads indeed limits cell type detection, we reprocessed the HPAP scRNA-seq datasets using Cell Ranger v7.0.1, which allows for the inclusion of intronic reads. After doing so, we successfully identified both c11 and c13 cells (Fig. 1d). We also evaluated where c11/c13 cells are positioned on the UMAP plot when using only exonic reads (Fig. 1f, left) versus both exonic and intronic reads (Fig. 1f, right). In the former case, both c11 and c13 cells are merged into the alpha cell cluster. However, when intronic reads are included, c11 cells form a distinct cluster, clearly separate from alpha cells. We further confirmed this observation in an independent dataset. Kang et al.^15^ had generated both scRNA-seq and snRNA-seq data from the same human islet samples; c11 and c13 cells were detectable in the snRNA-seq data and in scRNA-seq only when intronic reads were included (Fig. 1g). Thus, we identified two previously overlooked cell population, primarily missed due to the exclusion of intronic reads in prior scRNA-seq analyses.

### C11 and c13 cells are distinct from canonical endocrine cell types

Next, we analyzed the relationship of the newly identified cell clusters to endocrine cells. At the cluster level, we observed that c11 cells exhibit high expression of both glucagon and insulin (Fig. 2a). However, when examining the per-cell expression, we found a bipartite pattern: most c11 cells express high levels of glucagon, while a subset, located at the left edge of the branch, exhibited elevated insulin expression (Fig. 2b-c), comparable to that of mature beta cells. Notably, neither c11 nor c13 cells exhibit significant somatostatin expression (data not shown). The intriguing expression pattern of insulin and glucagon, combined with the importance of intronic reads in distinguishing this cell group, led us to investigate whether the c11/c13 cells represent residual fetal-like endocrine progenitors. To this end, we took advantage of a published scRNA-seq dataset profiling the fetal human pancreas^16^, which had identified a pan-endocrine progenitor population. However, when we mapped our multiome data onto this developing fetal human pancreas dataset, c11 and c13 cells aligned predominantly with alpha cells, not with endocrine progenitors (Supplementary Fig. 3a). Additionally, analysis with CytoTRACE2^17^ – a neural-net based model trained to learn gene expression patterns related to multipotency – suggested that c11/c13 cells are in a differentiated state with low developmental potential (Supplementary Fig. 3b), indicating that these cells are not remnant fetal endocrine progenitors.

**Figure 2:**
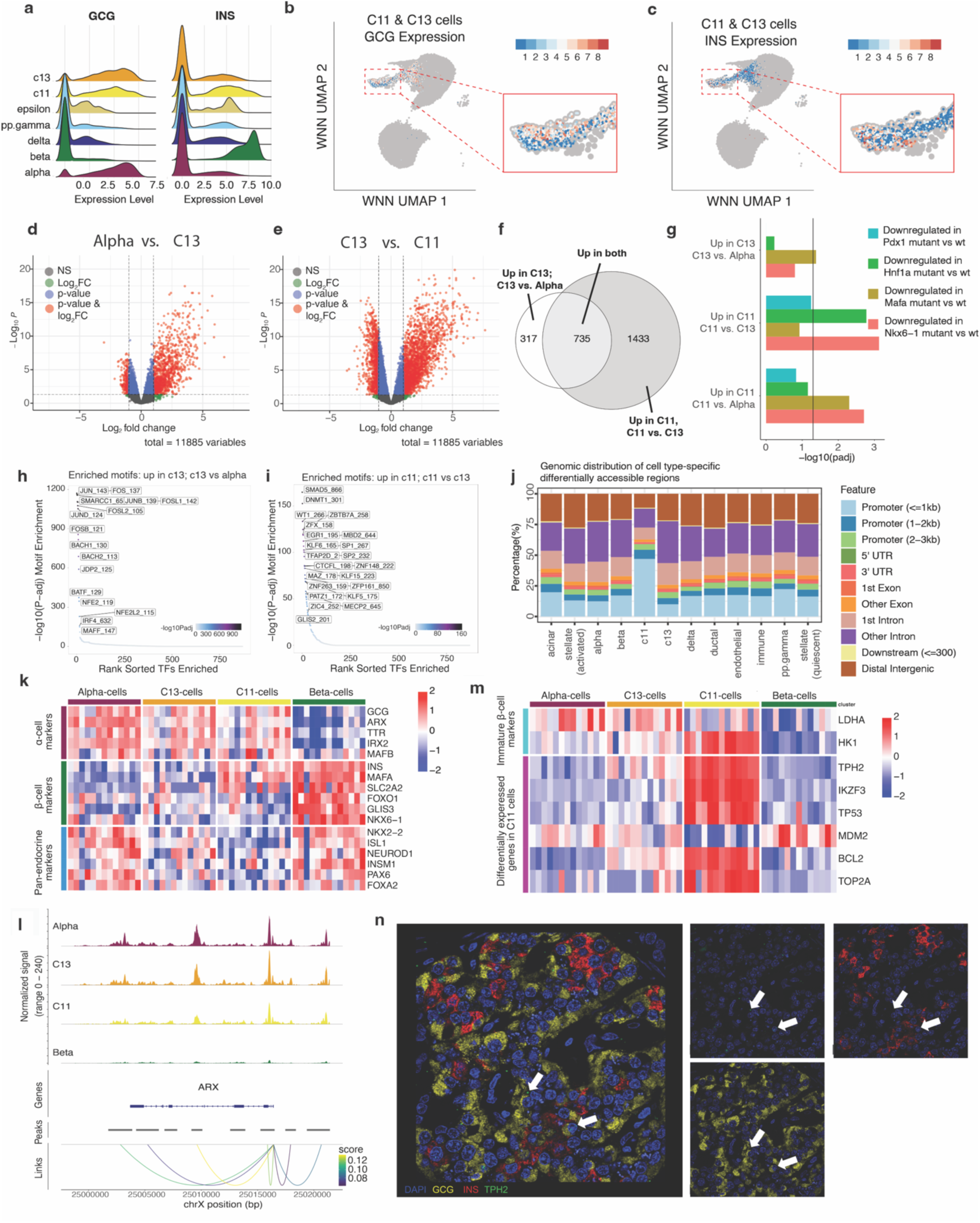
Transitional endocrine cells display both alpha and beta cell features. (a) Distribution of insulin (INS) and glucagon (GCG) expression across each endocrine cell type in the HPAP multiome dataset. (b-c) Expression of (b) INS and (c) GCG in transitional cells, overlaid on the WNN UMAP plot, with alpha-cells and beta-cells shown in gray. (d-e) Volcano plots of differential expression analysis: (d) comparison between alpha-cells and c13-cells, and (e) comparison between c13-cells and c11-cells. Differential expression was assessed using EdgeR^32^. (f) Venn diagram illustrating the overlap between genes upregulated in c13-cells compared to alpha-cells, and genes upregulated in c11-cells compared to c13-cells. (g) Enrichment analysis of differentially expressed genes in gene sets regulated by key beta-cell transcription factors, as determined by gene ablation studies in mice. Enrichment was calculated using a hypergeometric test with Benjamini-Hochberg correction. (h) Transcription factor binding motifs enriched in chromatin peaks with increased accessibility in c13-cells compared to alpha-cells. (i) Motifs enriched in chromatin peaks with increased accessibility in c11-cells compared to c13-cells. (j) Distribution of genomic annotations for peaks showing increased accessibility in one cell type versus all others. (k) Pseudo-bulk expression of alpha-cell, beta-cell, and pan-endocrine cell markers across selected cell types. Each row represents a gene, and each column represents a pseudo-bulk aggregation of cells from a single cell type and sample. Gene expression is shown as log-transformed counts-per-million, with rows scaled by gene. (l) Chromatin accessibility at the ARX locus, with peaks linked by peak-gene prediction analysis correlated with ARX expression. (m) Pseudo-bulk expression of selected markers upregulated in c11-cells. (n) RNA-Scope validation of TPH2 expression (green) in human islets, a marker differentially expressed in c11-cells. Note TPH2 expression in a subset of alpha cells. Alpha cells were labeled by immunofluorescence staining for glucagon (yellow), and beta cells were labeled by immunofluorescence staining for insulin (red).

It was recently observed that experimental cell dissociation, as required to produce single cell suspension from pancreatic islets, can alter gene expression profiles^18^. However, both c11 and c13 cells did not show a high percentage of reads from dissociation-induced genes (Supplementary Fig. 3c), making it highly unlikely that their discovery is an artifact of the experimental paradigm. Another possibility we considered was whether c11/c13 cells simply represent a special alpha-cell state, such that of cycling or senescent cells. However, c11/c13 cells do not express canonical markers of proliferation (MKI67, CDK1), senescence (CDKN1A/p21, CDKN2A/p16), or any gene signatures associated with apoptosis, cell cycle, senescence, or the senescence-associated secretory phenotype (SASP) (Supplementary Fig. 3d-e).

### Novel Population Represents Alpha-to-Beta Transition

Given the activation of the insulin gene in these alpha-cell related clusters, we hypothesized that c11 and c13 cells together represent a transitional state between alpha and beta cells. To explore this trajectory, we performed differential gene activity analysis using both gene expression profiles and chromatin accessibility data. In both data modalities, we compared c13 cells with either alpha cells or c11 cells. The differential gene expression analysis revealed a large set of genes upregulated in c13 cells, with a further subset activated in c11 cells (Fig. 2d-f). Many of these genes have been shown to be regulated by key transcription factors involved in beta-cell function through gene ablation studies in mice, including HNF1A, MAFA, and NKX6-1 (Fig. 2g). When analyzing chromatin accessibility, we focused on the transcription factors that could drive the alpha to c13 to c11 transition. From alpha cells to c13 cells, we identified motifs for proteins involved in stimulus-induced transcriptional activation, including FOSL1, JUN, and BACH1, which are part of the AP-1 complex^19^ (Fig. 2h). Comparing c11 cells with c13 cells, we found enriched motifs for SMAD5, MBD2, KLF5, and SP1, which bind to CpG-rich promoter regions^20–23^ (Fig. 2i). Consistent with this observation, newly accessible chromatin regions in c11 cells are highly enriched for promoter elements, specifically within 1kb of transcription start sites (TSS) (Fig. 2j), consistent with the *de novo* gene activation we observed in this cell cluster.

Next, we assessed the expression of key alpha-cell, beta-cell, and pan-endocrine markers (Fig. 2k). C11 and c13 cells displayed similar alpha-cell marker expression to alpha cells, but with progressively lower levels of the critical alpha-cell transcription factor ARX. In support of this finding, c11 cells also showed reduced chromatin accessibility at the *ARX* promoters and its enhancers (Fig. 2l). In contrast, beta-cell marker genes such as *INS*, *MAFA*, and *SLC2A2* (GLUT2) were upregulated in c11 cells (Fig. 2k). C11 cells exhibited lower expression of pan-endocrine cell markers (Fig. 2k), consistent with a more immature phenotype, and upregulation of *HK1*, a beta-cell disallowed gene typically expressed in immature beta-cells^24^(Fig. 2m). Finally, we identified Tryptophan hydroxylase 2 (TPH2) as marker for c11 cells in all donors analyzed (data not shown). RNA-Scope staining for TPH2 in human islets confirmed that a small subset of glucagon-positive cells expresses TPH2 (Fig. 2n). Other genes such as the Ikaros family zinc finger transcription factor IKZF3 and the DNA topoisomerase TOP2A, involved in chromatin remodeling, were also enriched in c11 cells (Fig. 2m). Notably, c11 cells exhibited higher levels of TP53 and lower levels of its antagonist MDM2, alongside with high expression of the anti-apoptotic gene BCL2, suggesting that these anti-DNA damage pathways may be activated due to large-scale chromosomal remodeling expected to occur during trans-differentiation.

To analyze this putative endocrine trans-differentiation process further, we performed trajectory analysis with ArchR, which produced a path from alpha to c11 via c13 cells (Fig. 3a-b). We then identified genes that exhibit coordinated changes in transcription factor gene expression and chromatin accessibility of their binding motifs along this trajectory (Fig. 3c-d). This analysis revealed broad downregulation of mature islet transcription factor genes such as *ISL1* and *HNF4A* and loss of accessibility at their associated binding sites (Fig. 3e). At the c13 to c11 stage transition, *CREB5*, which interacts with AP-1 complex subunits to activate transcription, becomes expressed. Another transcription factor gene showing gene activation and motif enrichment among accessible chromatin is the PAR bZIP factor *HLF*, recently suggested to be part of an islet gene signature^27^. Additionally, we observed the expected decrease in activity of the alpha cell master regulator *ARX* and an increase in both gene expression and gene activity scores for beta-cell transcription factors including *MAFA*, *PDX1*, and *NKX6-1*, especially at the tail end of the c11 cluster (Fig. 3f).

**Figure 3:**
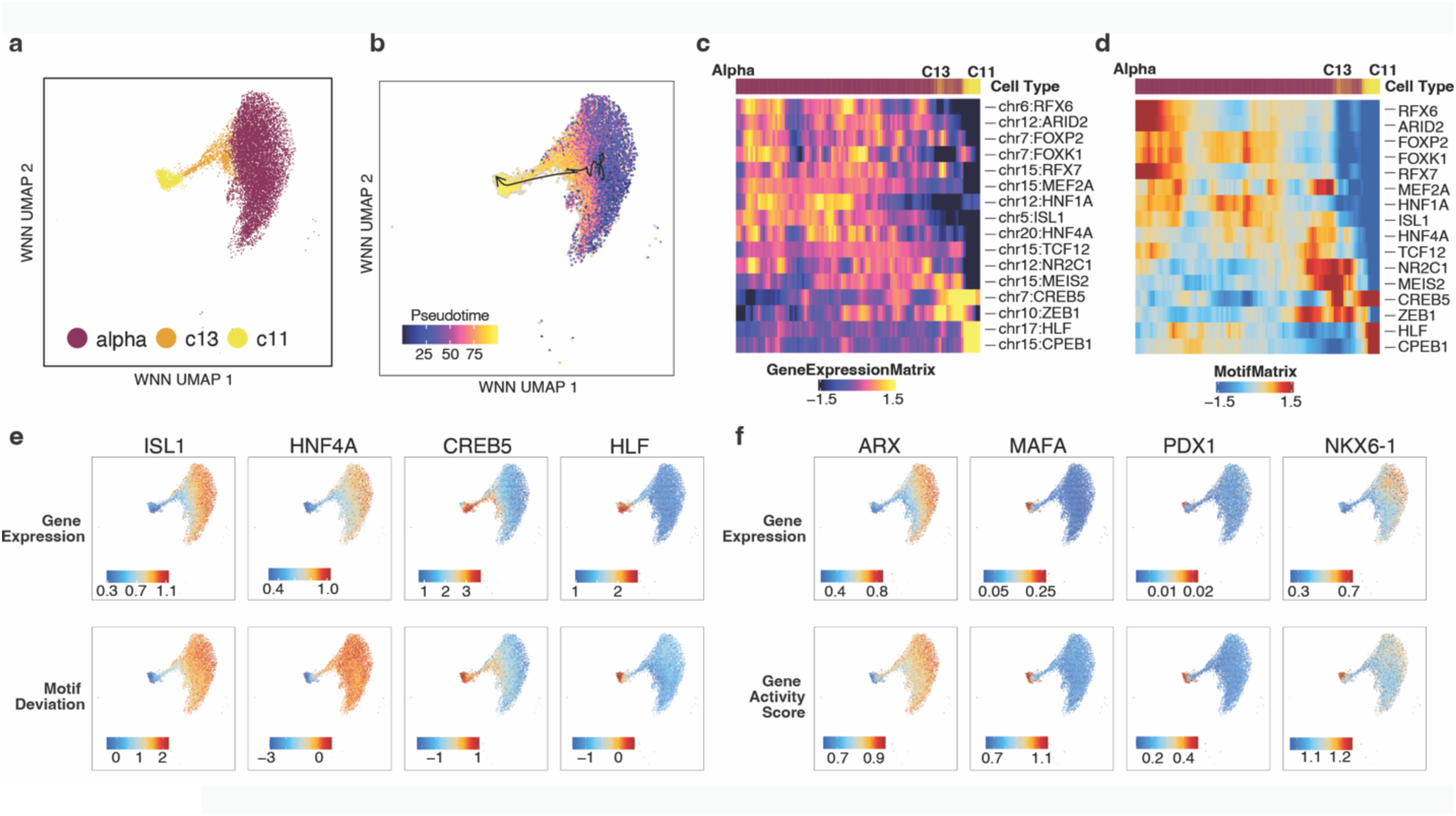
Pseudotime analysis suggests alpha cell conversion in human islets. (a) UMAP plot of c11-cells, c13-cells, and a subset of alpha-cells included in the trajectory analysis. (b) Inferred trajectory and pseudotime for the transition from alpha-cells to c13-cells to c11-cells, generated using ArchR^33^. (c-d) Heatmaps showing (c) gene expression and (d) chromVAR^34^ motif scores for transcription factors with correlated changes across RNA expression and their binding sites along the pseudotime trajectory. (e) Gene expression (top) and chromVAR motif scores (bottom) for key transcription factor in transitional cells. Both expression and motif scores are smoothed using MAGIC^35^ imputation. (f) Gene expression (top) and gene activity scores inferred from ATAC-seq data (bottom) for key transcription factors in alpha-cells and beta-cells. Both expression profiles are smoothed using MAGIC imputation.

### Increased Frequency of Transitional Cells in Islets from Donors with T2D

Next, we explored the possibility that transitional cells might be induced in the chronic hyperglycemic state of T2D. The HPAP cohort analyzed by single-cell modalities is well matched for body mass index (BMI) and age between non-diabetic donors and those with T2D (Fig.4a). When analyzing the abundance of transitional cells, we found a significantly higher proportion of transitional cells in islets from donors with T2D, compared to matched non-diabetic controls (Fig. 4a). Interestingly, this was the case for c13 cells but not c11 cells, suggesting that possibly the c13 to c11 transition presents a developmental “bottleneck”.

**Figure 4:**
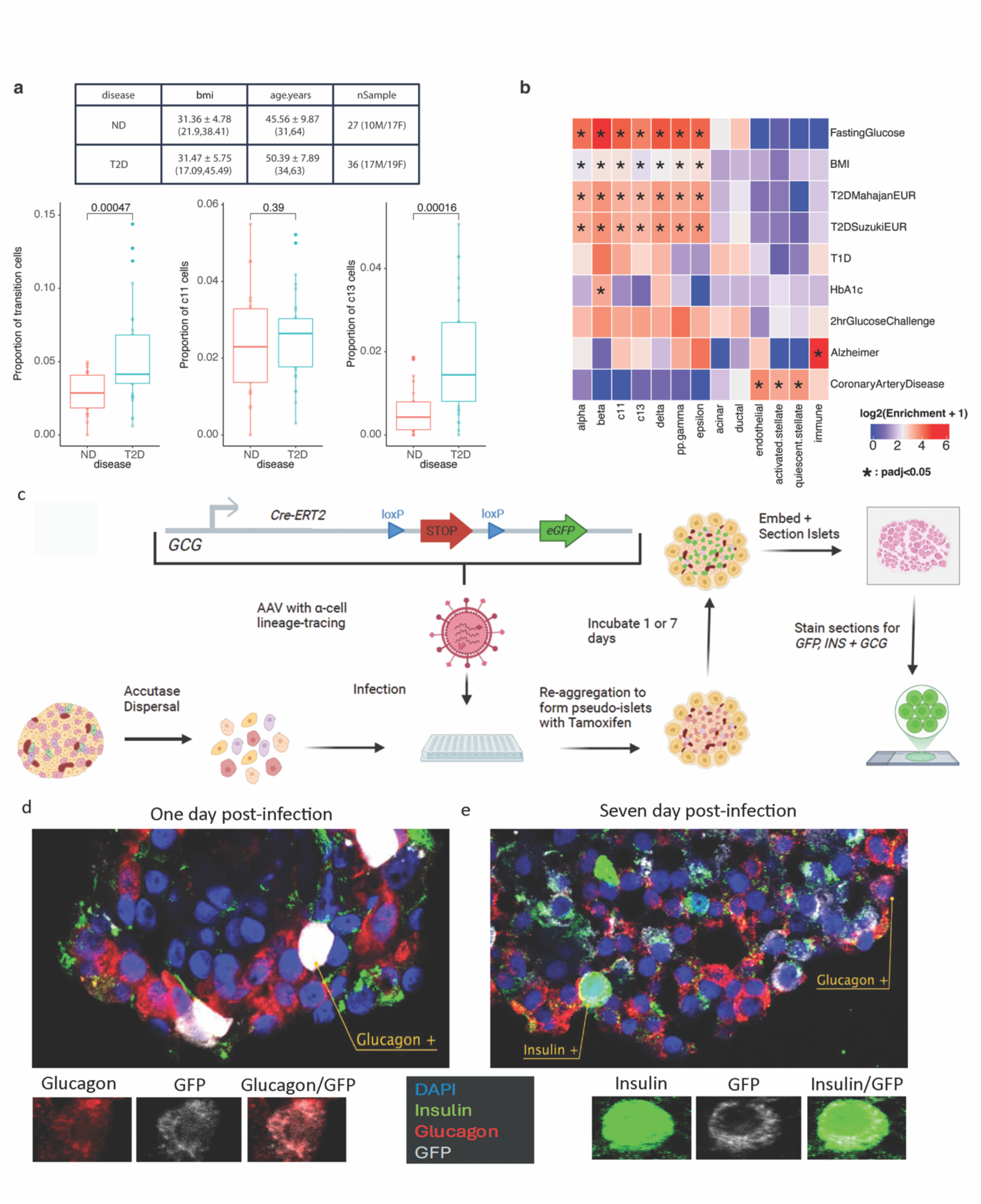
Abundance of transitional cells is increased in type 2 diabetes. (a) Comparison of transitional cell abundance in islets from donors with type 2 diabetes or non-diabetic controls. Top: Groups are matched by age, BMI, and sex. Bottom: The percentage represents the number of transitional cells relative to the total number of endocrine cells. (b) Enrichment of GWAS heritability in open chromatin regions across different cell types. The score represents the enrichment value calculated using stratified LD-score regression, normalized as log2(Enrichment + 1). Significant results with a Benjamini-Hochberg adjusted p-value < 0.05 are marked with an asterisk (*). (c) Schema depicting the procedure for genetic lineage tracing of alpha cells in primary human islets. (d) Representative image of islets incubated for one day post viral transduction and reaggregation before sectioning and staining. Immunofluorescence for glucagon (GCG: red), Insulin (Ins: green) and GFP protein (GFP: grey). Nuclei were counterstained with Hoechst dye (blue). Arrow points to a glucagon/GFP double positive cell. No dual insulin and GFP positive cells were observed at this time-point. (e) Representative image of islets incubated for one week post viral transduction and reaggregation before sectioning and staining. Color scheme as in (d). Arrows indicate glucagon/GFP double-positive and insulin/GFP double-positive cells, which are also shown in detail below.

We examined whether the chromatin regions accessible in c11 and c13 cells are enriched for the heritability of glycemic traits as measured by genome-wide association studies (GWAS). Using stratified LD-score regression (S-LDSC)^28,29^, we found that the chromatin regions accessible in c11 and c13 cells were enriched for the heritability of T2D, BMI, and fasting glucose, similar to what was seen for the mature endocrine cell types (Fig. 4b). Additionally, chromatin peaks that were preferentially accessible in transitional cells compared to non-endocrine cells were also enriched for fasting glucose and BMI heritability, supporting the idea that c11/c13 cells may be functionally important in metabolic regulation and diabetes etiology (Supplementary Fig. 4).

Finally, we employed genetic lineage tracing to ascertain if alpha to beta-cell transdifferentiation, as suggested by our single cell multiome analysis, could be observed in primary human islets. To this end, we designed an adeno-associated viral (AAV) vector to indelibly mark the alpha cell lineage using a tamoxifen-inducible, Cre-dependent and alpha cell-specific expression of green fluorescent protein (GFP) (Fig. 4c). Dispersed primary human islets were transduced with this virus, briefly treated with tamoxifen to induce Cre-mediated activation of the GFP reporter in alpha cells, and then cultured *in vitro* for one or seven days. After the one-day incubation, GFP expression was limited to alpha cells, as expected, confirming the cell type-specificity of the GCG promoter driven CreER transgene (Fig. 4d). However, after seven days of culture, GFP/insulin double positive cells were apparent (4e), indicative of spontaneous alpha to beta cell transdifferentiation. Thus, these data provide strong support for the notion of alpha to beta cell plasticity in the human endocrine pancreas.

## Discussion

In summary, we identified two previously undocumented cell populations in human islets, termed c11 and c13. We believe these cells may have been overlooked in prior single-cell studies because the default scRNA-seq pipeline excludes intronic reads. Given that c11 and c13 cells exhibit transitional phenotypes, capturing their actively transcribed gene profiles requires intronic reads to distinguish them clearly from other cell types. Of note, we conducted stringent quality control measures—including the removal of ambient RNA and filtering of doublets— minimizing the possibility of an artifact from the scRNA-seq workflow. Importantly, c11 and c13 cells are equally apparent in chromatin accessibility profiles; they separate distinctly from alpha cells in HPAP snATAC-seq data, even without mapping to the multiome reference. Unlike scRNA-seq, snATAC-seq is less susceptible to artificial doublets, as each cell’s chromatin profile is constrained to two copies per chromosome and does not suffer from the high background signal from highly expressed genes such as *INS* or *GCG* as the transcriptomic assays do. Additionally, c11 cells were identified in published datasets processed in independent laboratories, with different protocols, and we were able to validate expression of specific c11/c13 marker genes by RNAscope analysis on human pancreas sections. These findings give us strong confidence in the reliability of c11 and c13 cells as distinct populations. Finally, our genetic lineage tracing studies, performed on primary human islet cells, provide further evidence that glucagon-expression alpha cells can transition into insulin-expressing cells in humans without the necessity to ectopically overexpress beta cell transcription factors.

We propose that c11 and c13 cells represent at least a partial transdifferentiation of alpha into insulin-expressing cells. We hypothesize that this transition is driven by downregulation of the alpha-cell transcription factor ARX, leading to upregulation of beta-cell identity genes such as INS, MAFA, and SLC2A2. Notably, c11 cells express HK1, an immature beta-cell marker found in beta cells that have not yet reached full functional maturity^24^, suggesting that the transition progresses from alpha cells to beta cells via an immature beta-like intermediate, i.e. the c11 population. Further studies, such as Patch-seq^31^, which combines transcriptomic and electrophysiological measurements, could help clarify the functional state of c11 cells. Additionally, methods to sort c11/c13 cells would facilitate further profiling to better understand the mechanisms underlying this transdifferentiation process. Nevertheless, our data strongly support the notion that a subset of pancreatic alpha cells continuously transitions to insulin-positive beta cells in the human pancreas, a process that appears accelerated in individuals with T2D. Future work will be directed at identifying factors that might be used to accelerate this process in order to increase functional beta cell mass in people with diabetes.

## Methods

### Islet Procurement

Pancreatic islets were obtained through the HPAP consortium (RRID:SCR_016202; https://hpap.pmacs.upenn.edu), which is part of the Human Islet Research Network (https://hirnetwork.org). This study was approved by the University of Florida Institutional Review Board (IRB #201600029) and the United Network for Organ Sharing (UNOS). Informed consent was obtained from each donor’s legal representative prior to organ donation. The recovery and processing of organs followed established protocols^36^. Islets were cultured and dissociated into single cells as previously described^37^.

### Single Cell/Nucleus Molecular Profiling Assays

This study utilized three types of single-cell/nucleus molecular profiling assays: single-cell RNA sequencing (scRNA-seq), single-nucleus ATAC sequencing (snATAC-seq), and the multiome assay, which generates single-nucleus RNA-seq and ATAC-seq of the same cell. The scRNA-seq data presented here is an extension of a prior study^10^. Detailed protocols for each assay are available under the ’Workflow & Protocol’ section of HPAP’s PANC-DB website (https://hpap.pmacs.upenn.edu). As the HPAP consortium continuously acquires samples and conducts assays, there are ongoing updates to the kits and sequencing platforms used. Comprehensive metadata on the generation and sequencing of each sample is accessible on the PANC-DB website via the ‘Download All Metadata’ button on the ’Experiment Data Download’ page. Supplementary Table 1 summarizes the list of donors analyzed with each technology.

### Multiome Data Quantification and Quality Control

Cell Ranger ARC (v2.0.2) was used to demultiplex the sequencing output and generate two cell-by-feature matrices per sample, aligning reads to the GRCh38-2020-A-2.0.0 reference genome provided by 10x Genomics. Quality control was performed separately for each modality. For gene expression data, ambient RNA was removed using SoupX^38^ (v1.6.2). Doublets were detected and removed using functions from scDblFinder^39^ (v1.12.0). We then created a Seurat^40,41^ (v5.1.0) object, filtering out genes expressed in fewer than three cells. Low-quality cells were excluded based on the following thresholds: cells with fewer than 200 or more than 10,000 unique genes, cells with mitochondrial expression levels exceeding 30%, and cells with fewer than 1,000 or more than 100,000 total transcripts.

For chromatin accessibility data, we first generated a peak counts matrix for each sample. MACS2^42^ (v2.2.7.1) was used to call peaks, and the ‘FeatureMatrix’ function from Signac^43^ (v1.13.0) was employed to generate the cell-by-peak matrix. To remove doublets, we followed the ‘Doublet identification in single-cell ATAC-seq’ workflow^44^. Doublets were identified using Amulet^45^ which detects barcodes with more than 2 fragments overlapping the same genomic regions, and scDblFinder^39^ (v1.12.0) that filters cells by simulating artificial doublets. A combined p-value for doublet likelihood was calculated for each cell, and cells with a p-value < 0.05 were removed. Additional low-quality cells were filtered out based on nucleosome signal > 2, TSS enrichment < 1, peak counts fewer than 1,000 or exceeding 50,000, or when fragments mapping to ‘blacklist’ regions exceeded 0.05%. Only barcodes passing quality control in both modalities were retained for further analysis.

### Integration Across Samples for Gene Expression Data

To perform cell type annotation, we embedded all cells in a common space using both gene expression and chromatin accessibility profiles. We began by addressing batch effects within each modality, followed by integration across modalities. For gene expression data, we utilized Seurat v5’s streamlined integration method. Initially, gene expression data from all samples were combined, with donor IDs specified as layers. Each sample was normalized individually using scTransform^46^ (v0.4.1), and dimensionality reduction was performed via principal component analysis (PCA).

We tested three integration methods: (1) reciprocal PCA without reference samples (gexInt1), (2) reciprocal PCA with 9 reference samples (gexInt2), and (3) Harmony integration (gexInt3, Harmony v1.2.0). The integrated embedding from each method was passed to FindClusters (resolution = 0.5, method = igraph). Each resulting cluster was manually assessed and annotated. The nine high quality non-diabetic samples chosen as reference were: HPAP097, HPAP104, HPAP131, HPAP139, HPAP146, HPAP155, HPAP157, HPAP159, and HPAP160.

### Integration Across Samples for Chromatin Accessibility Data

For chromatin accessibility data, features were unified across samples by combining peaks identified individually using the ‘reduce’ function from GenomicRanges^47^ (v1.50.0), generating an updated cell-by-peak counts matrix per sample. Each sample was processed using the latent semantic indexing (LSI) workflow^48^. Two integration methods were applied: (1) reciprocal LSI projection using 9 non-diabetic samples as reference (atacInt1) and (2) Harmony integration (atacInt2). The integrated embeddings were then passed to ‘FindClusters’ (with parameters resolution = 0.5, method = igraph), and the resulting clusters were manually assessed and annotated.

### Integration Across Modalities

We compared the clustering results from each integration method, ultimately choosing to integrate gexInt2 (gene expression) and atacInt2 (chromatin accessibility) to represent each modality. The Weighted Nearest Neighbor (WNN) approach^40^ was used to integrate across modalities to obtain the final embeddings and clusters. After manual annotation, clusters showing mixed signals from multiple cell types were filtered out. The integration was then repeated for each modality separately, followed by the WNN integration, clustering, and manual annotation of the clusters based on marker gene expression.

### ScRNA-seq Data Quantification and Quality Control

For the HPAP scRNA-seq samples, we obtained counts with Cell Ranger (v7.1.0) using build GRCh38-2020-A as reference and with --include-introns set to true as per default. We used SoupX^38^ (v1.6.2) to remove ambient RNA contamination. For each sample, we then utilized Seurat v4.9.9.9041 to create a Seurat object (setting assay.version=“v3”, min.cells = 3, min.features = 200). After removing doublets with the Bioconductor package scDblFinder^39^ (v1.12.0), we applied the following filters: 200 < nFeature_RNA < 10000, percent.mt < 25, 500 < nCount_RNA < 100000 and then normalized the data with Seurat Log Normalization.

### ScRNA-seq Cell Type Annotation

We mapped scRNA-seq data to the multiome reference data using the supervised PCA (sPCA)^40^ approach. We first re-normalized the multiome gene expression data with Seurat Log Normalization. The top 2000 most variable genes were identified, and z-score standardization was applied. A sPCA transformation was learned between the normalized data and the weighted nearest neighbor graph from the WNN integration mentioned above. The learned sPCA transformation was then applied to each scRNA-seq sample, allowing the identification of transfer anchors between query and reference dataset. Using the identified anchors, scRNA-seq cells were projected to the reference UMAP space and a cell type label was predicted. We kept cells with prediction scores > 0.8 for the downstream analyses.

### SnATAC-seq Data Processing

Each HPAP snATAC-seq sample was processed using Cell Ranger ATAC (v2.0.0) with GRCh38-2020-A-2.0.0 as the reference genome. Using the fragment file and peak set provided by Cell Ranger, the peak count matrix was generated using the ‘FeatureMatrix’ function from Signac^43^ (v1.13.0). Low-quality cells were filtered out based on the following criteria: nucleosome signal > 2, TSS enrichment < 1, and peak counts fewer than 1,000 or exceeding 100,000.

### SnATAC-seq Cell Type Annotation

We annotated cell types in the snATAC-seq data by mapping it to the multiome reference using the supervised LSI (sLSI)^40^ approach. To avoid exceeding matrix size limitations in R, we downsampled the multiome dataset to 1,000 cells per donor. Peaks within the top 50th expression quantile were used to learn the sLSI transformation onto the WNN graph. For each snATAC-seq sample, a peak counts matrix was generated using peaks identified in the multiome data. We excluded samples with < 100 cells. The learned sLSI transformation was then applied to project each snATAC-seq dataset into the reference space, where cell type labels were predicted. Cells with a prediction score > 0.8 were retained for downstream analyses.

### ScRNA-seq Without Intronic Reads

We processed a subset of scRNA-seq data with Cell Ranger v5.0.1, which by default excludes intronic reads. To evaluate the effect of including intronic reads, we generated two UMAP plots of the same cells using an identical pipeline (Fig. 1f). The only difference between the plots is that the left plot was generated using a counts matrix from Cell Ranger v5.0.1, while the right plot used a counts matrix from Cell Ranger v7.0.1, which includes intronic reads. Cell barcodes present in both datasets after QC filtering were retained, followed by Seurat’s Log Normalization and rPCA integration (using 3,000 integration features, dims = 1:50), with four non-diabetic donors (HPAP052, HPAP103, HPAP104, HPAP105) set as the reference. Cells were labeled according to annotations from the **ScRNA-seq Cell Type Annotation** section.

### Mapping Kang *et al.* to HPAP multiome dataset^15^

We obtained the counts matrix for snRNA-seq and scRNA-seq from GSE217837. To generate a scRNA-seq counts matrix with intronic reads, we downloaded the SRA files associated with the scRNA-seq data. The SRA Toolkit (v3.0.7) was used to convert SRA files into FASTQ files. We then generated the counts matrix using Cell Ranger (v7.0.0) with --include-introns set to true as per default and used GRCh38-2020-A as reference. For scNA-seq (exon-only) and snRNA-seq, we applied Seurat’s Log Normalization and projected each islet donor sample onto the multiome reference data using sPCA. For scRNA-seq (exon + intron), we applied the same filters as described in the original manuscript: nFeature_RNA > 250, percent.mt < 20, nCount_RNA > 500, log10GenesPerUMI > 0.8, followed by Log Normalization and reference mapping with sPCA. We retained cells with prediction score > 0.8.

### Mapping to the Ma *et al.* Human Fetal Pancreas Dataset^16^

We obtained scRNA-seq counts matrix and metadata file from the OMIX database with accession number OMIX001616. We used cells from post conception week 7 -11. We processed the data using the standard Seurat pipeline with Log Normalization. Reciprocal PCA was used to map each of the HPAP scRNA-seq and multiomeRNA onto the human fetal pancreas dataset. The cell type label released by the original authors^16^ was used for annotating the HPAP cells. Cells with prediction score > 0.8 were retained. We also used another reference mapping method, SingleR^49^ (v1.8.1). We converted Seurat objects to SingleCellExperiment^50^ (v1.16.0) objects, computed log-normalized expression values, and ran SingleR (using de.method=”wilcox”). Each cell was annotated with the label that had the highest score and we retained cells with prediction score > 0.5.

### Stemness Prediction

We predicted stemness in each cell of the multiome dataset using CytoTRACE2^17^ (v1.0.0). For each donor, we created a Seurat object containing the multiome data and passed it to the ‘cytotrace2’ function (specifying slot_type = ’counts’ and species = ’human’). This generated a predicted potency score for each cell, reflecting stemness. We then calculated the median potency score for each cell type within each donor and visualized these values in a box plot (Supplementary Fig. 3b) to compare stemness across cell types.

### Dissociation Test

We obtained a list of dissociation-induced genes from Supplementary Table 5 of van den Brink *et al*.^18^ . Gene names were standardized to uppercase, and only those present in the HPAP single-cell datasets were included in the analysis. To quantify the effects of dissociation, we calculated the percentage of reads corresponding to dissociation-induced genes using the “PercentageFeatureSet” function from Seurat. Cells among the top 6% expression quantile of the dissociation expression are considered as dissociation affected.

### Differential Expression Analysis

Differential expression test was performed using two packages, EdgeR^32^ (v3.36.0), under R 4.1.3. Pseudo-bulk samples with >=30 number of cells and library size >= 50,000 number of reads were analyzed. To identify differentially expressed genes in transitional cells, we set up the EdgeR object with donors that have pseudo-bulk samples passing the threshold above for all four cell types of interest: alpha-cells, beta-cells, c13-cells and c11-cells. We excluded long non-coding genes from the input gene expression matrix. We filtered out genes with low expression by using ‘filterByExpr’ (with min.count=5 and min.total.count=15). For this analysis, the glmQLFTest was employed with “∼celltype + donor” as the design. Then we performed pairwise analysis between every combination by specifying a contrast matrix. We reported differentially expressed genes between c13 cells vs. alpha-cells, or c11 cells, respectively. Genes with adjusted p-value < 0.05 and log_2_(foldchange) > 1 were retained as differentially expressed genes.

### Gene Set Enrichment Test

We performed gene set enrichment test using the hypergeometric test, followed by multiple testing correction using the Benjamini-Hochberg procedure^51^. We obtained gene sets representing downstream targets of key beta-cell transcription factors, specifically, genes downregulated in beta-cell specific gene ablation of Hnf1a^52^, Pdx1^53^, Nkx6-1^54^, and Mafa^55^. Additionally, we sourced the following gene sets from MSigDB^56^: HALLMARK_APOPTOSIS, KEGG_APOPTOSIS, WP_APOPTOSIS, GOBP_CELLULAR_SENESCENCE, GOBP_CELL_CYCLE, GOBP_CELL_CYCLE_DNA_REPLICATION,

HALLMARK_PANCREAS_BETA_CELLS. The senescence associated secretory phenotype (SASP) gene list (referred to as SAUL_SASP in this study) was obtained from Saul et al^57^.

### ATAC-seq analyses

We used ArchR^33^ (v1.0.1) for the downstream analyses of the ATAC-seq data. Arrow files were generated using the fragment files outputted from Cell Ranger ARC. We retained cells present after both ArchR and Seurat/Signac processing (see **Multiome Data Quantification and Quality Control**). We filtered out five samples that have median number of fragments per cell < 8,000. Cell type specific peaks were called by first merging cells of the same type using ‘addGroupCoverages’ with parameters: minCells=100, maxCells=500, minReplicates=3, maxReplicates=8. Peaks were called per cell type using MACS2^42^ and merged using the iterative overlap procedure previously described^58^. We identified differentially accessible regions (DARs) for each cell type by running “getMarkerFeatures”, which performs Wilcoxon test to compare cells of one cell type versus background cells from the other cell types while also matching TSS enrichment and sequencing depth. ChIPSeeker^59,60^ was used to annotate peaks to genomic regions. To identify the drivers of the transition between alpha to c11 cells, we identified DARs in c13 vs. alpha-cells and c11 vs. alpha-cells. DARs with false discovery rate (FDR) ≤ 0.1 and log2 fold-change ≥ 0.5 were passed to motif enrichment where motifs were obtained from the CIS-BP database^61^.

### Trajectory Analysis

We used ArchR functions to order c11, c13, and alpha-cells in pseudo-time. Cells were processed with the iterativeLSI procedure^33^, and Harmony^62^ was employed to integrate cells across samples. With the Harmony-adjusted embedding, we learned a trajectory based on a prespecified order of alpha-cells to c13-cells to c11-cells. Given the large population of alpha-cells not involved in the transition, we filtered out cells with a pseudo-time > 50 and reran the trajectory inference workflow. We calculated gene activity scores from ATAC-seq data using addGeneScoreMatrix. Motif deviation score was calculated using ChromVAR^34^ per cell. Trajectory plots were smoothed using weights imputed from MAGIC^35^ (by calling ArchR::addImputeWeights function with reducedDims=’Harmony’ and k=8).

### Abundance Test

We calculated the proportion of c11, c13, and transitional cells (c11+c13) relative to the total endocrine cells per donor. For donors with multiome data, we used the proportion from the multiome dataset as the measure of abundance. For donors with both scRNA-seq and snATAC-seq data, we averaged the two proportions, while for donors with only one dataset, that measurement was used as-is. We only used samples with > 100 endocrine cells. For the T2D vs. non-diabetic (ND) comparison, ND donors were selected as controls based on the absence of all four autoantibodies and age > 30. Grubbs’ test was applied iteratively to identify and remove outliers, repeating until the highest value in each group was no longer significant. Each group had 1 to 4 values removed. A likelihood ratio test was used to compare a full model with a reduced model that excluded the ‘disease’ variable in the linear regression. Gender, age, body mass index (BMI), and race were included as covariates in the regression model.

### GWAS enrichment

We use stratified LD-score regression (S-LDSC)^28,29^ (v1.0.1) to calculate the heritability enrichment of various traits in the called ATAC-seq peaks of pancreatic cell types. For this analysis we obtained summary statistics of GWAS for T2D^63,64^, T1D^65^, HbA1c^66^, fasting insulin (FI)^66^, fasting glucose (FG)^66^, 2-hour glucose challenge^66^, body mass index (BMI)^67^, coronary artery disease (CAD)^68^, and Alzheimer’s disease^69^. Where necessary, we converted summary statistics to GRCh38 using the University of California Santa Cruz (UCSC) Liftover tool^70^. For each trait/cell-type combination, we used the S-LDSC pipeline to estimate enrichment values relative to a background consisting of the baseline LD model (v2.2) and merged peaks across all the pancreatic cell-type clusters. We used the 1000 Genomes Phase III European-ancestry individuals genotyped in GRCh38 as an LD reference for this analysis and treated the ATAC-seq peaks as binary annotations. Following the recommendations of the S-LDSC authors, we bounded the unbiased enrichment estimates output by S-LDSC to values ≥ 0. All estimates < 0 were assigned a *p*-value of 1 for the purposes of determining enrichment significance, and we corrected for multiple-testing using the Benjamini-Hochberg procedure^51^.

### RNAscope validation for c11 markers

RNAscope was performed on formalin-fixed-paraffin-embedded (FFPE) slides obtained through HPAP including samples from T2D and ND donors. Slides were baked at 60℃ for approximately 1 hour in an ACD Hybez Hybridization oven (ACD, 241000). Deparaffinization was done by immersing slides in fresh 100% Xylene twice for 5 minutes each, followed by immersion in 100% Ethanol twice for 2 minutes each. Due to variable fixation quality across tissues from different donors, protease treatment, and target retrieval steps were skipped for optimal results. Following deparaffinization, slides were dried at 60℃ for 5 minutes. A hydrophobic barrier was created around the tissue sections using an Immedge Pen (Vector Laboratories, H-4000), and allowed to dry completely before proceeding.

Slides were briefly washed in UltraPure Nuclease Free Distilled Water (Thermo Fisher, 10977015). RNA probes and wash buffer were pre-warmed at 40℃ for 10 minutes, then left to cool down to room temperature. For hybridization, a single probe was used per tissue section. C1 probes were used at 1X concentration, while C2 and C3 probes were diluted in probe diluent at 1:50 (ACD, 300041). Probes were applied to cover tissue sections, and slides were incubated at 40℃ for 2 hours.

Wash Buffer (1X) was prepared from the (50X) stock provided by the manufacturer (ACD, 310091) using UltraPure Nuclease Free Distilled Water. Following Hybridization, slides were washed in an ACD EZ-Batch Wash Tray twice for 2 minutes at room temperature (ACD, 321717). Amplification steps in AMP1, AMP2, AMP3 were done consecutively at 40℃ according to the manufacturer’s recommended protocol, with two washes for 2 minutes at room temperature in between each incubation.

Fluorophores were diluted 1:1500 in the TSA Buffer provided by the manufacturer. After removing excess liquid, HRP was applied to cover each tissue section entirely. The HRP reagent was matched to the channel of the mRNA probe. For instance, HRP-C3 was used for the Hs-FGF18-C3 mRNA probe. Slides were incubated at 40℃ for 15 minutes. Slides were washed twice for 2 minutes in fresh wash buffer and incubated at 40℃ for 30 minutes in the fluorophore mix. Slides were washed twice more in fresh wash buffer and treated with HRP blocker for 15 minutes at 40℃. Finally, slides were washed twice for 2 minutes before proceeding to the protein immunofluorescent staining.

We performed immunofluorescence staining on the same slides undergoing RNAscope. Immediately following the RNAscope assay, slides were washed twice in PBST for 2 minutes each. They were then incubated at room temperature for 30 minutes in CasBlock (Thermofischer, 008120). Primary antibodies were diluted to a working concentration of 1:100 in CasBlock and applied to slides for a 1 hour incubation at room temperature. Primary antibodies include anti-mouse Glucagon (Boster Bio, MA1047), anti-goat CD99 (R&D Systems, AF3968-SP), and anti-rabbit Insulin (Cell Signaling Technology, 3014S). After incubation in primary antibodies, slides were washed twice in PBST for 2 minutes. Secondary antibodies were diluted in PBS-1% BSA at 1:300 and applied to slides for 1 hour at room temperature. Secondary antibodies include Cy2 AffiniPure Donkey Anti-Mouse (Jackson Labs, AB_2340825), Cy3 AffiniPure Donkey Anti-Goat (Jackson Labs, AB_2340411), and Donkey anti-Rabbit Alexa Fluor 790 (Thermofischer, A11374). Slides were washed twice in PBS for 5 minutes, and excess liquid was removed. Tissue sections were covered in DAPI reagent (provided in the RNAscope kit). Excess DAPI was removed by flicking the slides, then the slides were then immediately mounted using ProLong Gold Antifade Mounting medium (Thermo Fisher, P36930).

| Reagents | Catalog # |
| --- | --- |
| RNAscope Multiplex Fluorescent Reagent Kit v2 | ACD, 323100 |
| RNAscope Probe Diluent | ACD, 300041 |
| ImmEdge Hydrophobic Barrier Pen | Vector Laboratories, H-4000 |
| ACD HybEZ Hybridization oven | ACD, 241000 |
| RNAscope Probe - Hs-FGF18-C3 - Homo sapiens fibroblast growth mRNA | ACD, 444711-C3 |
| ACD EZ-Batch Wash Tray | ACD, 321717 |
| HybEZ Humidifying paper | ACD, 310015 |
| RNAscope Multiplex TSA Buffer | ACD, 322809 |
| CAS-Block Histochemical Reagent | Thermo Fisher, 008120 |
| Corning Cell Culture Buffers: Dulbecco's Phosphate-Buffered Salt Solution 1X | Corning life Sciences, MT21031CV |
| UltraPure DNase/RNase-Free Distilled Water | Thermo Fisher, 10977015 |
| UltraPure BSA | Thermo Fisher, AM2616 |
| Tween 20, 100% Nonionic Detergent | Bio-Rad, 1706531 |
| ProLong™ Gold Antifade Mounting medium | Thermo Fisher, P36930 |
| Opal 690 Reagent Pack | Akoya Biosciences, SKU FP1497001KT |
| Cy2 AffiniPure Donkey Anti-Mouse IgM | Jackson Labs, AB_2340825 |
| Cy3 AffiniPure Donkey Anti-Goat IgG (H+L) | Jackson Labs, AB_2340411 |
| Donkey anti-Rabbit IgG (H+L), Alexa Fluor 790 | Thermo Fisher, A11374 |
| Anti-Glucagon Antibody (Monoclonal, K79bB10) | Boster Bio, MA1047 |
| Insulin (C27C9) Rabbit mAb | Cell Signaling Technology, 3014S |

### Genetic lineage tracing in human pseudo-islets

96 well agarose gels were prepared for incubation of dissociated cells to reaggregate them into pseudo-islets. Briefly, 2g of Ultrapure Agarose (Invitrogen: 16500500) were dissolved in 10mLs of 0.9% saline in an Erlenmeyer flask and microwaved until completely dissolved. Then, 500uL of agarose was pipetted into 3D Petri Dish® micro-mold spheroids (Millipore Sigma: Z764043) to form 96-well gels in which re-aggregation of cells to pseudo-islets can take place. After the agar solidified, the gel was flexed out of the mold and into a 12 well plate. The well was then filled with 2mLs of prewarmed PIM(S)® media (Prodo labs cat:123456789) to equilibrate/prepare the wells and kept at 37°C in an incubator until needed.

Non-diabetic donor islets were obtained through the Human Pancreas Analysis Program (HPAP) and incubated in PIM(S)® media overnight. Islets were poured into non-treated 100mm petri dishes (Sigma: P5731) and counted under a dissecting microscope before transfer to 10mLs of PIM(S)® media in a 15mL Falcon tube (Fisher Scientific: 14-959-53A). The islets were then centrifuged at 200g at room temperature for 2 minutes before addition of 500μL of pre-warmed Accutase (Sigma: C-41310) per 500 islets. The islets were then disassociated in a 37°C water bath for 10 minutes before quenching with a 1:1 volume of 20% FBS (Hyclone:SH30910003)/DPBS (gibco: 14190). The dissociated islet cells were spun down at 300g for 3 minutes before washing with 1mL of PIM(S)® and repeating the centrifugation to finally resuspend the cells in 1mL of PIM(S)® media. Islet cells were then counted in a Countess 3 Automated Cell Counter using 20uL of a 1:1 dilution of cells + 0.4% Tryptan Blue (Corning: 25-900-CI). 200k cells were then pipetted into Eppendorf™ DNA LoBind™ Tubes (Eppendorf™ 022431021) at 2,000 cells/μL to be infected with α-cell lineage tracing AAV (Vectorbuilder: VB240827-1652cqu). The virus was thawed from -80°C at room temperature and gently vortexed prior to adding to cells. 8μL virus was added to the corresponding tubes and the cells were transduced with the virus for one hour with trituration at 10-minute intervals. The transduced cells and virus were then moved to 96 well plates (Thermo Scientific: 249952) and incubated overnight at 37°C in a tissue culture incubator (5% O_2_), after which they were moved from their wells into Lobind tubes and the wells washed twice with 75 μL of warmed PIM(S)® to collect all the cells. The cells were then centrifuged at 200g for 5 minutes and resuspended in 100 μL of PIM(S)® containing tamoxifen to wash out any remaining virus and allow activation of Cre-recombinase in alpha cells. Finally, the cells were pipetted dropwise into the prepared 96-well agarose molds, placed in 24 well plates and incubated for 72 hours at 37°C in a tissue culture incubator (5% O_2_) to allow the cells to aggregate and form pseudo-islets. After 72 hours, the agar gels were inverted to allow the pseudo-islets to be collected in 2mLs of PIM(S)® in the 24 well plates and incubated for 1 day or 7 days before processing for immunofluorescence.

Pseudo-islets were collected into a 1.5mL Lobind tube and washed with 1mL of PBS twice to remove any debris or dead cells with gentle centrifugation at 200g for 1 minute to pellet the pseudoislets between each wash. The supernatant was then removed and the islets fixed in 500 μL of formalin at room temperature for 30 minutes. The fixative was then aspirated and the islets washed twice with 1mL of PBS to remove residual fixative. Concurrently, Histogel (Cancer Diagnostics: SKU: HG10144) was melted at 70°C and 20 μL of Affi-Gel Blue Gel beads (Biorad: 1537301) were washed with PBS and resuspended after a quick spin on a tabletop centrifuge in 30 μL of PBS. The tube containing the islets was then aspirated to remove any remaining supernatant before addition of the beads/PBS mixture, which were spun together before removal of any remaining PBS. Finally, 20 μL of warm Histogel was added to the islet/bead mixture and mixed gently to avoid bubbles before pipetting onto a glass slide in a circular disc. This process was repeated twice with the Histogel pipetted progressively further on the outside for a total of 60 μL to ensure collection of all islets. The slide was then placed on a flat surface of wet ice for 10 minutes to allow the gel to solidify before sliding it into a 20mL scintillation vial containing PBS. The gel was washed 4 times with PBS for 30 minutes each prior to the dehydration steps. Using progressively more hydrophobic ethanol solutions (30% for 2 hours, 50% for 2 hours, 70% overnight, 95% for 3 hours twice, 100% for 1 hour twice), the samples were prepared for standard paraffin embedding to form blocks which were then sectioned at a thickness of 5 μm. Subsequently, the sections were stained for immunofluorescence as described above using antibodies for glucagon (Proteintech: 15954-1-AP), insulin (Proteintech: 66198-1-Ig) and *GFP* (Novusbio: NB100-1770).

### Data Availability and Code Availability

The raw and processed single-cell data generated during and during the current study will be available on the HPAP PANC-DB website (https://hpap.pmacs.upenn.edu) upon acceptance. The analysis pipeline will be available via the GitHub repository.

## Acknowledgements

We thank Dr. Golnaz Vahedi for valuable comments on the manuscript and Dr. Agnes Kloechendler for sharing the beta-cell transcription factor mutant gene sets. We also extend our gratitude to Dr. Avital Swisa for the fruitful discussions in the early phase of this study. We thank the current and past members of the HPAP team for their efforts in generating the data. Our heartfelt gratitude goes to the pancreas donor and their families for their invaluable donation, which made this study possible. This manuscript used data acquired from the Human Pancreas Analysis Program (HPAP-RRID:SCR_016202) Database (https://hpap.pmacs.upenn.edu/), a Human Islet Research Network (RRID:SCR_014393) consortium supported by NIDDK grants UC4-DK-112217, U01-DK-123594, UC4-DK-112232, and U01-DK-123716.

## Author Contributions

M.Y.Y.L. performed most computational and statistical data analysis; E.M., and J.S. performed pre-processing for some of the data; M.C. and B.V performed the LD Score regression. O.G. designed and performed the genetic lineage tracing study; O.G and H.E. performed experimental validations; H.D., D.L., T.D., C.L., and A.N. generated the single-cell data as part of the HPAP Consortium. M.Y.Y.L. and K.H.K wrote and all authors edited the manuscript; and K.H.K and M.L. supervised the study.

## Supplemental Information Supplementary Figures

**SF1:**
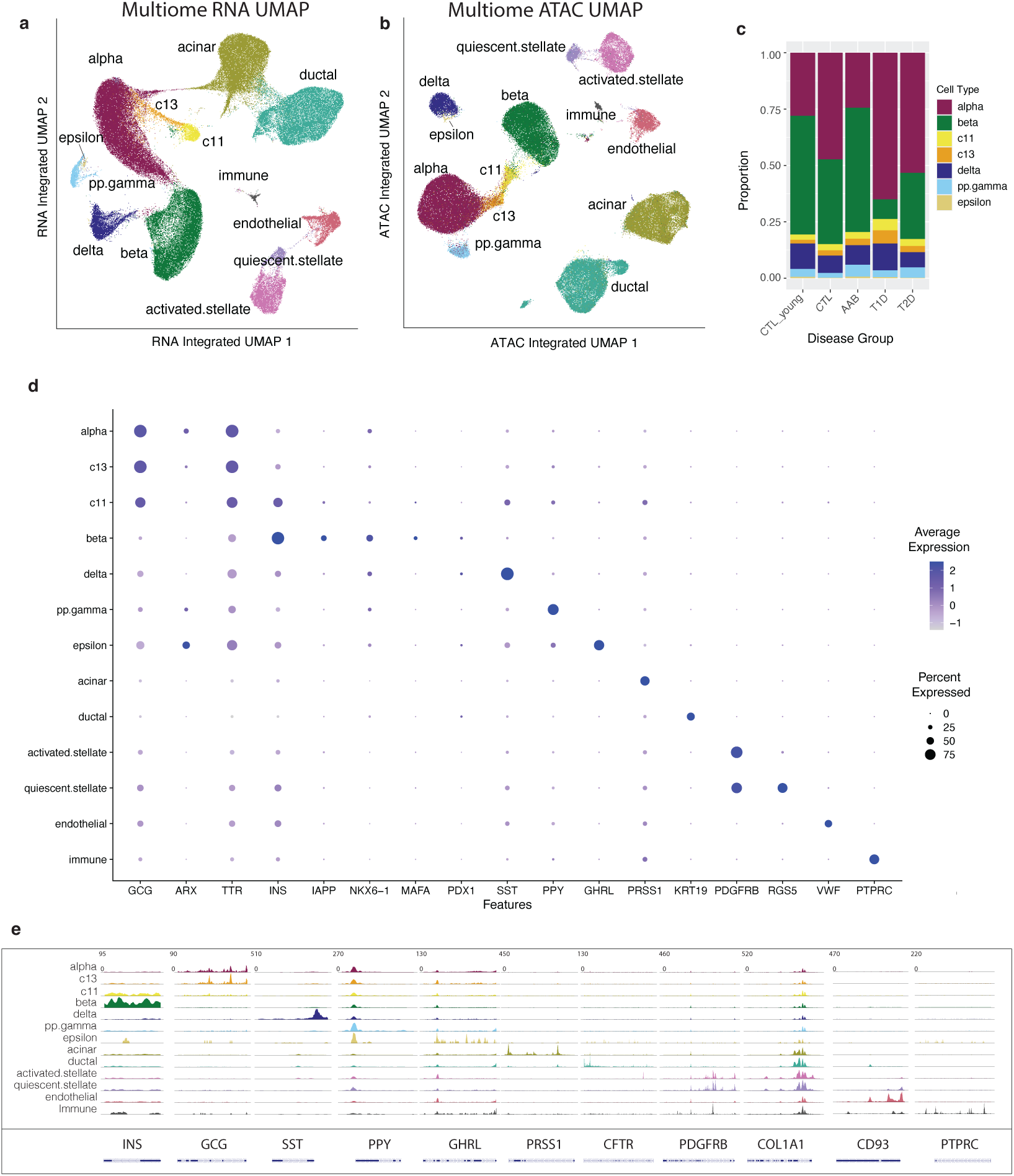
Cell type annotation result for the HPAP multiome data. (a) Visualization of cell type annotation of the multiome data, shown in UMAP space generated using the gene expression profile alone. (b) Cell type annotation shown in UMAP space generated using chromatin accessibility profile alone. (c) Bar plot showing proportion of each islet cell type across disease groups. C11 and c13 cells are present in all disease groups. (d) Dot plot showing marker gene expression in each cell type. (e) Coverage plot showing accessibility at marker genes, each row represents the accessibility profile of all cells in that cell type merged.

**SF2:**
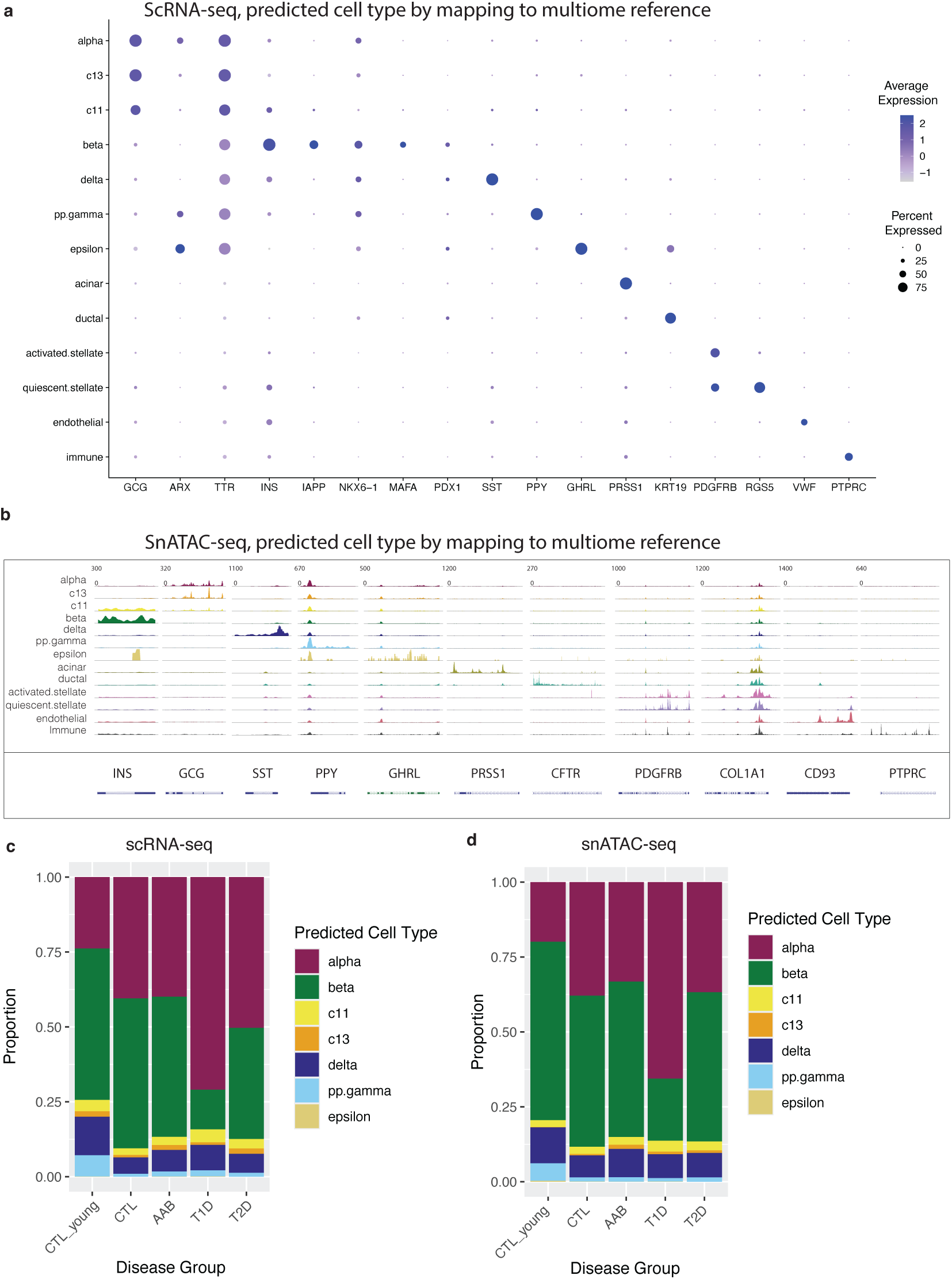
Cell type annotation result for HPAP scRNA-seq and snATAC-seq, after mapping to the HPAP multiome data. (a) Visualization of average expression of each marker gene across cell types. (b) Visualization of aggregated accessibility profile at each marker gene across cell type. (c) Bar plot showing proportion of each islet cell type across disease groups, for HPAP scRNA-seq. (d) Bar plot showing proportion of each islet cell type across disease groups, for HPAP snATAC-seq.

**SF3:**
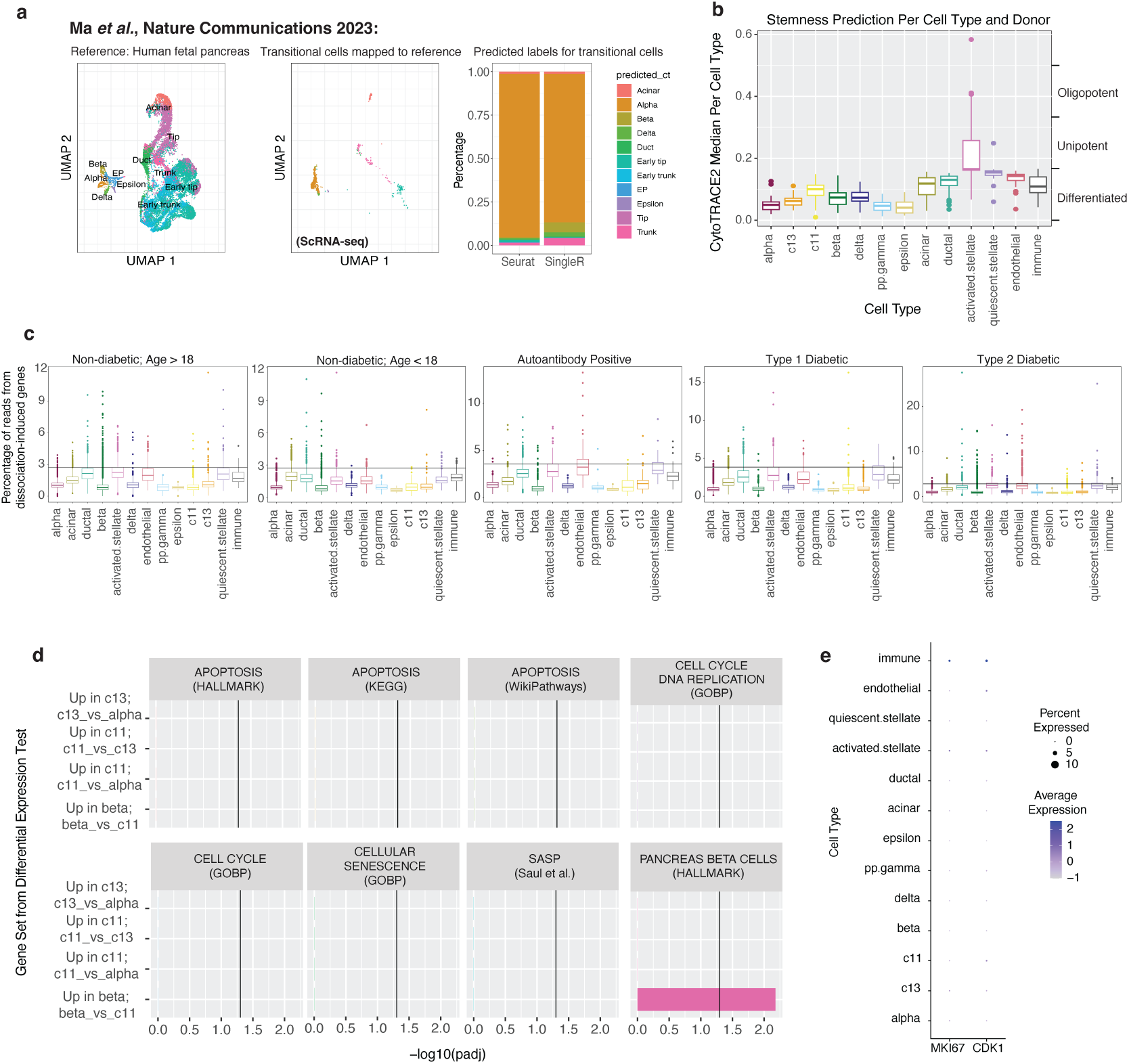
Investigation of transition cell identity. (a) UMAP plot of the human fetal pancreas reference dataset from Ma et al., Nature Communications 2023 (left), the UMAP plot of the HPAP scRNA-seq dataset, mapped onto the reference space (middle), bar-plot showing the distribution of predicted labels for transition cells in scRNA-seq dataset (right), most cells are predicted to be alpha-cells by both Seurat label transfer and SingleR prediction algorithm. (b) Prediction result from CytoTRACE2, grouped by cell type. The score shows the stemness of each cell and c11/c13 cells are annotated as ‘differentiated’ state. (v) Percentage of reads from dissociation-induced genes for each cell type, separated by disease groups. Dissociation-induced genes are from van den Brink *et al*., Nature Methods 2017. The horizontal line represents the 94% quantile per disease group. Cells with dissociation score > 94% quantile value are dissociated-affect cells. Transition cells (c11 and c13-cells) do not have high dissociation score. (d) Result of overrepresentation test, checking whether genes upregulated in c11/c13 cells are enriched for apoptosis, cell cycle or senescence signatures. (e) Dot-plot showing genes cell replication markers such as MKI67 and CDK1.

**SF4:**
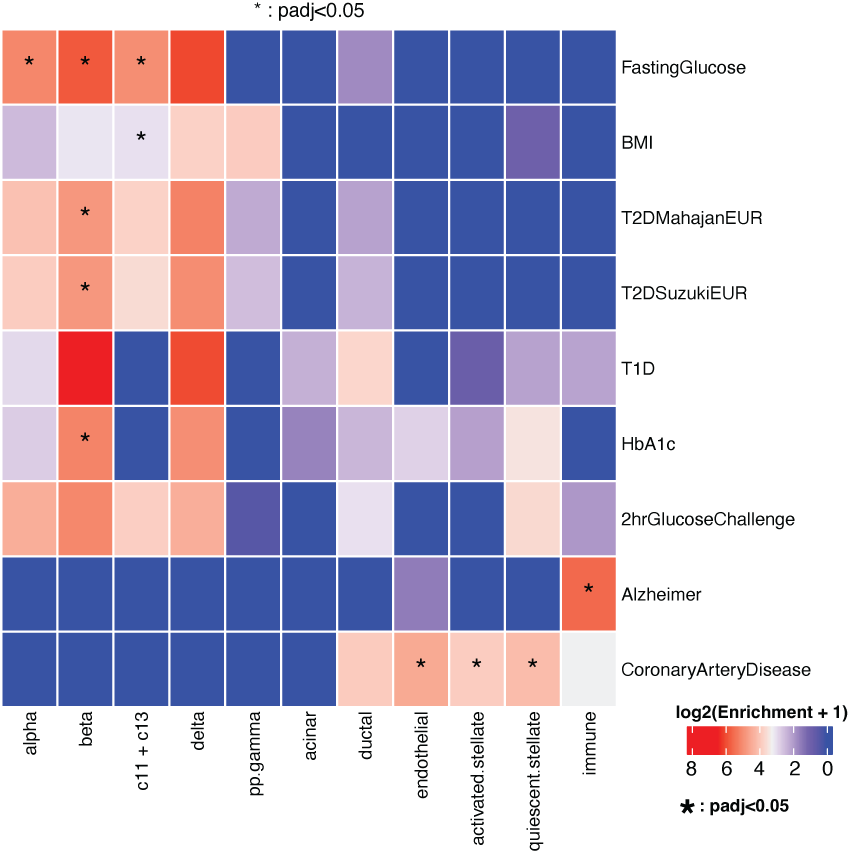
Enrichment of GWAS signals in differentially accessible regions across different cell types. The score represents the enrichment value calculated using stratified LD-score regression, normalized as log2(Enrichment + 1). Significant results with a Benjamini-Hochberg adjusted p-value < 0.05 are marked with an asterisk (*).

